# Investigating Statistical Power of Differential Abundance Studies

**DOI:** 10.1101/2024.06.07.597956

**Authors:** Michael Agronah, Benjamin M. Bolker

**Affiliations:** Department of Mathematics and Statistics, McMaster University, 1280 Main Street West, L8S 4K1, Ontario, Canada; Departments of Mathematics & Statistics and of Biology, McMaster University, 1280 Main Street West, L8S 4K1, Ontario, Canada

**Keywords:** Statistical Power, Effect Size, Differential Abundance, Microbiome

## Abstract

Identifying microbial taxa that differ in abundance between groups (control/treatment, healthy/diseased, etc.) is important for both basic and applied science. As in all scientific research, microbiome studies must have good statistical power to detect taxa with substantially different abundance between treatments; low power leads to poor precision and biased estimates via the “winner’s curse”. Several studies have raised concerns about low power in microbiome studies. In this study, we investigate statistical power in differential abundance analysis. In particular, we present a novel approach for estimating the statistical power to detect effects at the level of individual taxa as a function of effect size (fold change) and mean abundance. We analysed seven real case-control microbiome datasets and developed a novel method for simulating microbiome data. We illustrate how power varies with effect size and mean abundance; our results suggest that typical differential abundance studies are underpowered for detecting changes in individual taxon.

## Introduction

Identifying taxa that show differential abundance between groups holds great potential for clinical applications. For example, a study aimed at assessing the effects of a dietary intervention on microbial composition might analyse the abundance of different microbial taxa between a control group on a standard diet and a treatment group on a gut-health-promoting regimen.

Power analysis helps in minimizing false positives, thereby contributing to the trustworthiness of significant results obtained in studies. The power of a statistical test is the probability of successfully rejecting the null hypothesis given a particular effect size (Cohen, 2013). Power is determined by the sample size, effect size and the significance threshold (or “alpha level”), as well as methodological factors such as experimental design, number of groups, statistical procedure and model, type of response variable and fraction of missing data.

Power analysis enables researchers to detect meaningful effects and allocate resources efficiently; it aids the reliability and reproducibility of research findings. The primary goal of power analysis is to ensure that a research study has the sensitivity required to detect meaningful effects (Cohen, 2013). Underpowered studies are likely to miss biologically meaningful effects and are more prone to type II errors, which can lead researchers to neglect differences that could be biologically interesting (Goodman and Berlin, 1994). Even if a low-powered study finds statistically significant results, the estimated effect size will be imprecise (Goodman and Berlin, 1994). Low power together with a statistical significance filter (for example, only reporting effects with a *p*-value *<* 0.05) can lead to overestimation of the true effect (“magnitude”, or type M, error) or an incorrect estimate of the direction of an effect (“sign”, or type S, error) (Gelman and Carlin, 2014).

Microbiome researchers typically focus on three main types of analysis: (1) *analysis of univariate summaries*: reducing the data from each microbiome sample to a single value, such as alpha diversity, and comparing the distribution of these values between groups, (2) *community-wide analyses* using tests such as PERMANOVA or the Dirichlet-multinomial model to distinguish overall differences in communities, and (3) taxon-by-taxon or *differential abundance* analyses: identifying taxa with biological meaningfully differences between groups. Existing studies on power analysis have focused either on studies comparing univariate (alpha diversity) measures or studies comparing changes in overall microbiome composition between groups (Xia et al., 2018). For example, La Rosa et al. (2012) developed a reparameterized Dirichlet Multinomial model and a method for estimating the power to detect changes in overall microbial compositions between groups. Kelly et al. proposed a framework for estimating power in PERMANOVA.

To our knowledge, no methods exist for power analysis for differential abundance studies. In practice, every taxon in a microbial community has a different mean abundance and a different effect size (as is typical, we use fold change between groups as effect size in this paper), leading to a different statistical power to detect differences in every taxon.

Except for relatively simple analyses, conducting power analysis requires data simulation. Simulating an entire microbial community is challenging because it requires estimating appropriate community-wide distributions for mean abundances and effect sizes of taxa.

Power estimates in a differential abundance study depends on the abundance of individual taxa. For example, effect sizes of taxa with high abundance in both control and treatment groups are more likely to be detected compared to effect sizes of taxa that are rare in both groups. Unlike univariate power analysis where one can specify a single value for effect size and power, in a taxon-by-taxon power analysis there are multiple values of effect size and power; one for each taxon. This typically means hundreds or thousands of effect sizes and power values.

Several studies have raised concerns about low power in microbiome studies (Brüssow, 2020; Kers and Saccenti, 2021). For example, Kers and Saccenti showed that microbiome studies comparing alpha and beta diversities (PERMANOVA) between groups might be underpowered. The goal of this study is to investigate the issue of potential low power to detect effect size of individual taxon within a differential microbiome study. We develop a novel method for simulating microbial communities and estimating power at the level of an individual taxon. Our framework estimates power for each taxon as a function of effect size and mean abundance of an individual taxon. Using our framework, researchers can estimate the range of power in their studies and power for specific taxa and the expected number of significant taxa that will be detected in their study.

## Methods

### Data Collection and Proccessing

We examined seven microbiome datasets obtained from the European Nucleotide Archive (EBA) (Leinonen et al., 2010) and the National Center for Biotechnology Information (NCBI) (Sayers et al., 2021) in order to choose appropriate distributions for mean abundances and fold changes.

We performed a search using the query terms *“autism[All Fields] AND 16S[All Fields]”* and *“autism[All Fields] AND 16S[All Fields] AND Fecal[All Fields]”* on November 6, 2021. The search resulted in 10 datasets with accession numbers PRJNA168470, PRJNA3550, PRJNA453621, PRJEB45948, PRJNA644763, PRJNA589343, PRJNA687773, PRJNA578223, PRJNA624252 and PRJNA642975. Each data included a “treatment” group of children with autism spectrum disorder and a “control” group of neurotypical children.

To prepare the datasets for downstream analysis, we removed adaptors and primer sequences using the `cutadapt` function. We then processed the trimmed sequences into Amplicon Sequence Variant (ASV) data using the `Dada2` (Callahan et al., 2016) pipeline, which involved various steps such as filtering and trimming, error estimation, denoising, merging paired reads, and removing chimeras (Chen et al., 2020). Implementations of the `c` utadapt function in a bash script and the `Dada2` workflow in an `R` script are provided in the supplementary material. Three of the datasets (PRJNA578223, PRJNA624252, and PRJNA642975) had very low count abundances. Pre-filtering performed to remove low mean abundances resulted in the exclusion of the majority of taxa from these datasets. Consequently, these datasets were excluded, leaving seven datasets for our analysis.

### Model Description

The negative binomial model, a standard approach for analysing microbiome count data, is implemented in the `DEseq2` (Love et al., 2014) and `edgeR` (Robinson et al., 2010) R packages which are both widely used in microbiome analysis. The model is described as follows: Let *K*_*ij*_ denote the count data for the *i*^*th*^ taxon in the *j*^*th*^ sample. Then *K*_*ij*_ follows a negative binomial distribution:

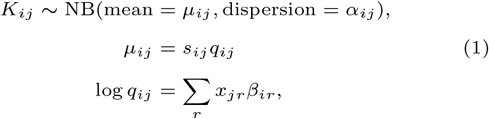

where *µ*_*ij*_ and *s*_*ij*_ are the mean abundances and normalization constants respectively. *q*_*ij*_ is the expected mean abundance of a given taxon in a sample prior to normalization. We assume the normalization constants *s*_*ij*_ are constant within a sample and the dispersion parameter is constant for a given taxon. Thus, *α*_*ij*_ = *α*_*i*_ and *s*_*ij*_ = *s*_*j*_. The estimated coefficients 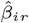 are estimates of the effect sizes and *x*_*jr*_ are the covariates. The relationship between the variance of counts and the dispersion is defined by var 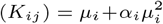. In this study, the estimating procedure implemented in the `DESeq2` package (Love et al., 2014; Anders and Huber, 2010) in `R` is used for estimating 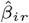 and 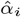

### Pre-filtering low abundant taxa and effect size shrinkage

Taxa with low abundance tend to exhibit high variability, which can pose challenges in detecting significant differences between groups. As is routinely done in differential abundance analysis, we filtered rare taxa, retaining only those taxa that had an abundance of five or more reads in at least three samples (Xia et al., 2018; Love et al., 2014). Rare taxa often lead to implausibly large fold change estimates. To tackle this problem, we used a shrinkage functionality in the `DESeq2` package, which shrinks large fold change estimates for low-abundance taxa towards zero.

### Method for simulating microbiome data

In order to simulate microbiome data for power calculation, we estimated community-wide distributions of mean abundance and fold change. The following sections describe our method for fitting distributions to the mean abundance and fold change of a particular data set, and the method for simulating microbiome data from the negative binomial model described in equation 1.

#### Modelling overall log mean abundance and log fold change

We modelled log mean abundance (that is, log of the arithmetic mean abundance from both control and treatment groups) as a finite mixture of Gaussian distributions. To determine the optimal number of components, we used a parametric bootstrap approach to sequentially test mixtures with 1 to 5 components. We used the implementation of the parametric bootstrap in the `mixtool` R package (Benaglia et al., 2010). For each successive pair of components (*k* and *k*+1 components), we conducted a parametric bootstrap by generating 100 bootstrap samples from the null model (the model with *k* components) and fitted both the null and alternate model (i.e., the model with *k* + 1 components) for each bootstrap sample to calculate a distribution of the likelihood ratio statistic under the null hypothesis. This statistic is used to test the null hypothesis of a *k* component fit against the alternative hypothesis of a *k* + 1 component fit across different mixture models. A *p*-value (with a 5% significance threshold) is used as a decision rule for selecting the optimal number of components. Once the *p*-value is exceeds the significance threshold, the testing terminates and the null model for the test where the procedure terminates is chosen as the number of components (Benaglia et al., 2010).

We also modelled log fold change as a finite mixture of Gaussian distributions. Fold change is typically related to mean abundance (Love et al., 2014). Thus, we modelled the mean and standard deviation parameters of the individual Gaussian components as functions of log mean abundance. In order to determine an appropriate way to model log fold change as a function of log mean abundance, we examined the relationship between log mean abundance and log fold change for each data. Figure 1 shows the relationship between log mean abundance and log fold change for three of the microbiome datasets. The smooth line representing the mean of log fold change as a function of log mean abundance appears to follow a linear trend. We modelled the mean parameter for each Gaussian component as a linear function of log mean abundance. Consequently, the overall mean of the mixture distribution is also a linear function of log mean abundance.

**Fig. 1.**
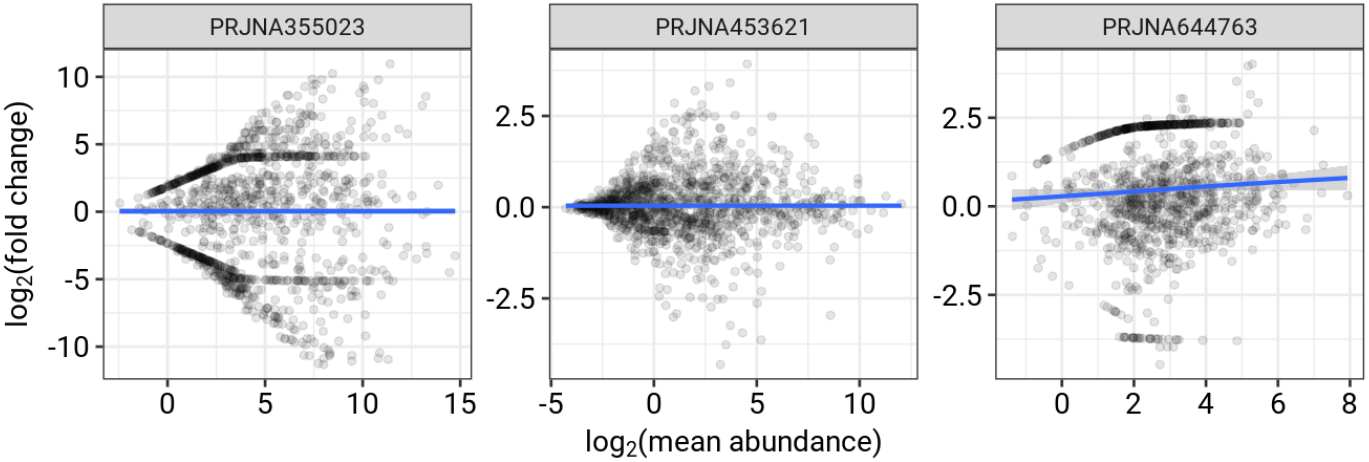
Relationship between log fold changes and log mean abundance for three typical datasets. The unusual features in the plot (concentrations of points along symmetric curves above and below zero) in the first two panels) correspond to taxa with zero counts across all subjects in either the control or the treatment group.

Upon examining variations of log fold change around the smooth line, we observed either a linear or quadratic trend (see scale-location plot in Figure 8 under supplementary materials). We therefore modelled the variance of each Gaussian component as both linear and quadratic functions of log mean abundance. We compared Gaussian mixtures with 1-5 components. For a given model (Gaussian mixture model with aspecified number of components), we modelled the variance parameter of all components either by a linear or quadratic function. We selected the model that yielded the minimum Akaike Information Criteria (AIC) value across all the fitted components.

The model of log fold change as a function of log mean abundance is described as follows:

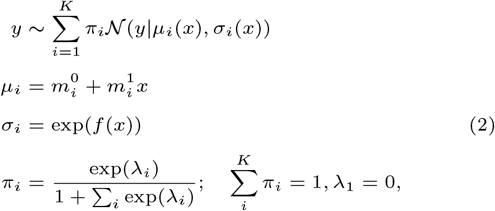

where *K* is the number of Gaussian components. *µ*_*i*_ and *σ*_*i*_ are the mean and standard deviation of the *i*^*th*^ component, conditional on the log mean abundance *x. y* is the log fold change. The function *f* denotes a linear or quadratic function of log mean abundance used to model the standard deviation parameter and *π*_*i*_ is the mixture probability with parameter *λ*_*i*_.

#### Modelling dispersion

We used the `DESeq2` package to estimate dispersion for the negative binomial model. Dispersion typically varies based on count abundance, with rarer taxa exhibiting higher dispersion (Love et al., 2014). To accommodate this variability and to simulate dispersion for subsequent power analyses, we used a nonlinear function of mean abundance to model the dispersion estimates, as implemented in the `DESeq2` package:

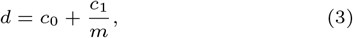

where *d* and *m* denote the scaled dispersion and mean abundance respectively. The term *c*_0_ represents the asymptotic dispersion level for high abundance taxa, and *c*_1_ captures additional dispersion variability.

Figure 9 in the supplementary material shows the spread of the dispersion estimates from the `DESeq2` package for each data. These estimates were extremely high (for example, dispersion values of 150 and 200). Using these dispersion estimates, we simulated count data from a negative binomial model with mean abundance from the microbiome dataset and log fold change estimates from the `DESeq2` package. The variability in the coefficient of variation of taxa abundances computed from the dispersion estimates were notably greater than observed in the actual dataset (see Figure 10 under supplementary materials). We therefore scaled the dispersion to align the coefficients of variation from the simulated data more closely with those from the observed datasets. A scaling factor of 0.3 was chosen based on trial and error to match the simulated and observed distributions of coefficient of variation for each microbiome data.

#### Data simulation procedure

The following steps outline our procedure for simulating microbiome count data.

- **Simulate overall log mean abundance:** For each data set, we simulated log mean abundance from the fitted Gaussian mixture distribution.
- **Simulate log fold changes:** Using the simulated log mean abundance, we simulated log fold change from the fitted Gaussian mixture distributions (equation 2).
- **Predict dispersion values:** Next, we predicted dispersion values as a function of the simulated mean abundance from the fitted non-linear function (equation 3).
- **Calculate per-group mean abundance:** We then calculated mean abundance for control and treatment groups using the simulated mean abundance and the simulated log fold change.
- **Simulate count data:** Using the calculated mean abundance for control and treatment groups and the predicted dispersion values, we simulated count abundances from the negative binomial model (equation 1).

We compared results from our simulation method with two existing methods for simulating microbiome data implemented in the `HMP` (La Rosa et al., 2012) and the `metaSPARSim` (Patuzzi et al., 2024) packages in R. The `HMP` package simulates microbiome data from a Dirichlet multinomial model and the `metaSPARSim` package simulates microbiome data from a Multivariate hypergeometric model.

### Method for estimating statistical power

We estimate statistical power as the probability of rejecting the null hypothesis (that the mean abundance in control and treatment group are the same for a given taxon). We used the `DESeq2` package to compute *p*-values for each taxon, using the Benjamini and Hochberg method for false discovery rate correction. The event that a given taxa is significantly different between groups is a Bernoulli trial. To estimate statistical power for various combinations of log mean abundance and log fold change, we fitted a shape-constrained generalized additive model (GAM) (Pya and Wood, 2015). The model predicting fold change as a function of log mean abundance is as follows

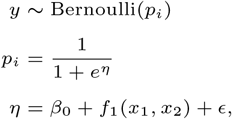

where *y* is a binary value (with 1 indicating that the *p*-value was below a critical value and 0 otherwise). We used the default critical value of 0.1 in the `DESeq2` package. *p*_*i*_ is the statistical power for taxon *i. β*_0_ and *ϵ* are the intercept and error terms respectively and the predictors *x*_1_ and *x*_2_ are the log mean abundance and log fold change respectively. The function *f*_1_ is a two-dimensional smoothing surfaces with basis generated by the tensor product smooth of log mean abundance and log fold change.

Power and fold change are positively correlated. Additionally, effect size of taxa with high abundance are more likely to be detected, hence having higher power, than rare taxa. To account for these relationships, we constrained the function *f*_1_ to be a monotonically increasing function of both log mean abundance and log fold change.

### Expected number of taxa with significant difference between groups

Consider a differential abundance study involving *n* taxa, each associated with power (probability of being significantly different between groups) *p*_*i*_. Whether taxon *i* differs significantly between groups or not in a particular analysis is a Bernoulli random variable with a success probability *p*_*i*_. Therefore, the expected number of significant taxa can be computed as the sum of the expected number of successes in *n* Bernoulli trials:

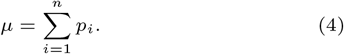

Equation 4 can be divided and multiplied by *n* to obtain

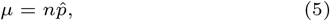

where 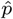 is the average statistical power across all taxa. Equation 5 states that the expected number of significant taxa in a differential abundance study is the product of the number of taxa (*n*) and the average statistical power for all taxa 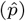.

## Results

Figure 2 and 3 compare the mean and variance distributions of taxa for three of the microbiome datasets with distributions from simulations made using the `HMP`, `metaSPARSim`, and our simulation method. For each microbiome dataset, we simulated count data using the same number of taxa and number of samples per group as the real data set.

**Fig. 2.**
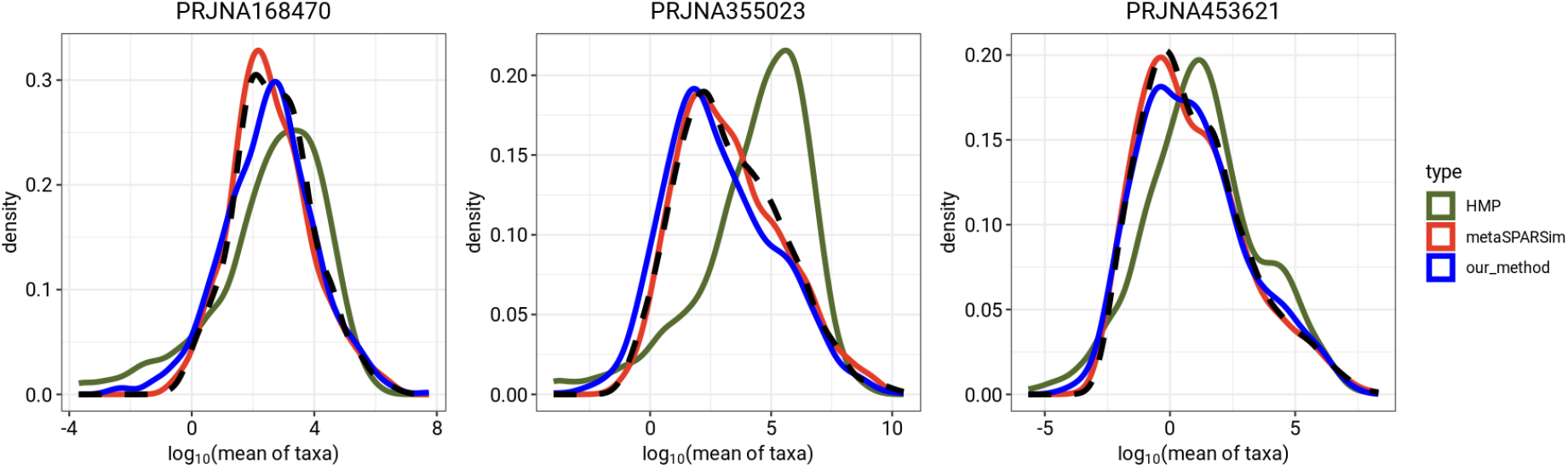
Comparison of distributions of mean abundance of taxa between observed data and simulations generated from `HMP`, `metaSPARSim` and our method. Black dash lines represent the distribution of mean abundance of taxa for the microbiome dataset

**Fig. 3.**
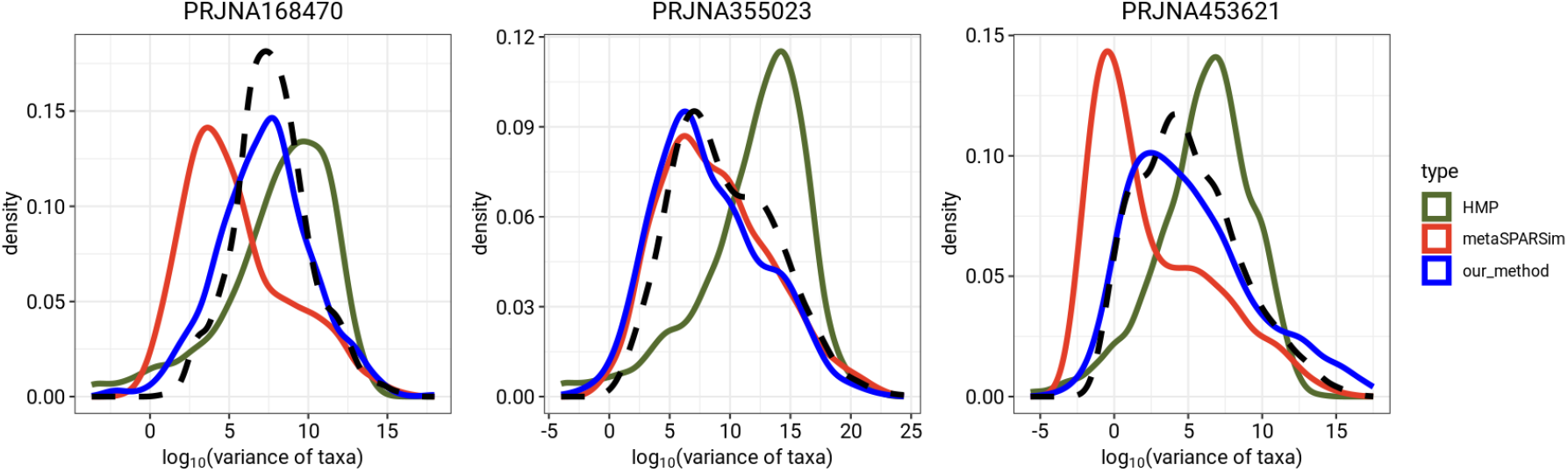
Comparison of the distributions of variance of taxa between observed data and simulation generated from `HMP`, `metaSPARSim` and our method. Black dash lines represent the distribution of variance of taxa for the microbiome dataset

Figure 4 shows the statistical power for all combinations of log mean abundance and log fold change for each data. Red points indicate simulated taxa with significant log fold change (that is, adjusted *p*-value *<* 0.1); black points show simulated taxa where we failed to reject the null hypothesis (adjusted *p*-value *>* 0.1). Contour lines show the predicted statistical power for various combinations of overall log mean abundance and log fold change. Figure 5 show the relationship between statistical power and the number of samples per group (30, 50, 70, 90, 110, 130, 150, 170 and 190 samples per group) for different log fold changes (2, 3 and 4). As expected, statistical power increases with increasing number of samples per group and increasing log fold change (Figure 5), although the power levels vary hugely across dataset.

**Fig. 4.**
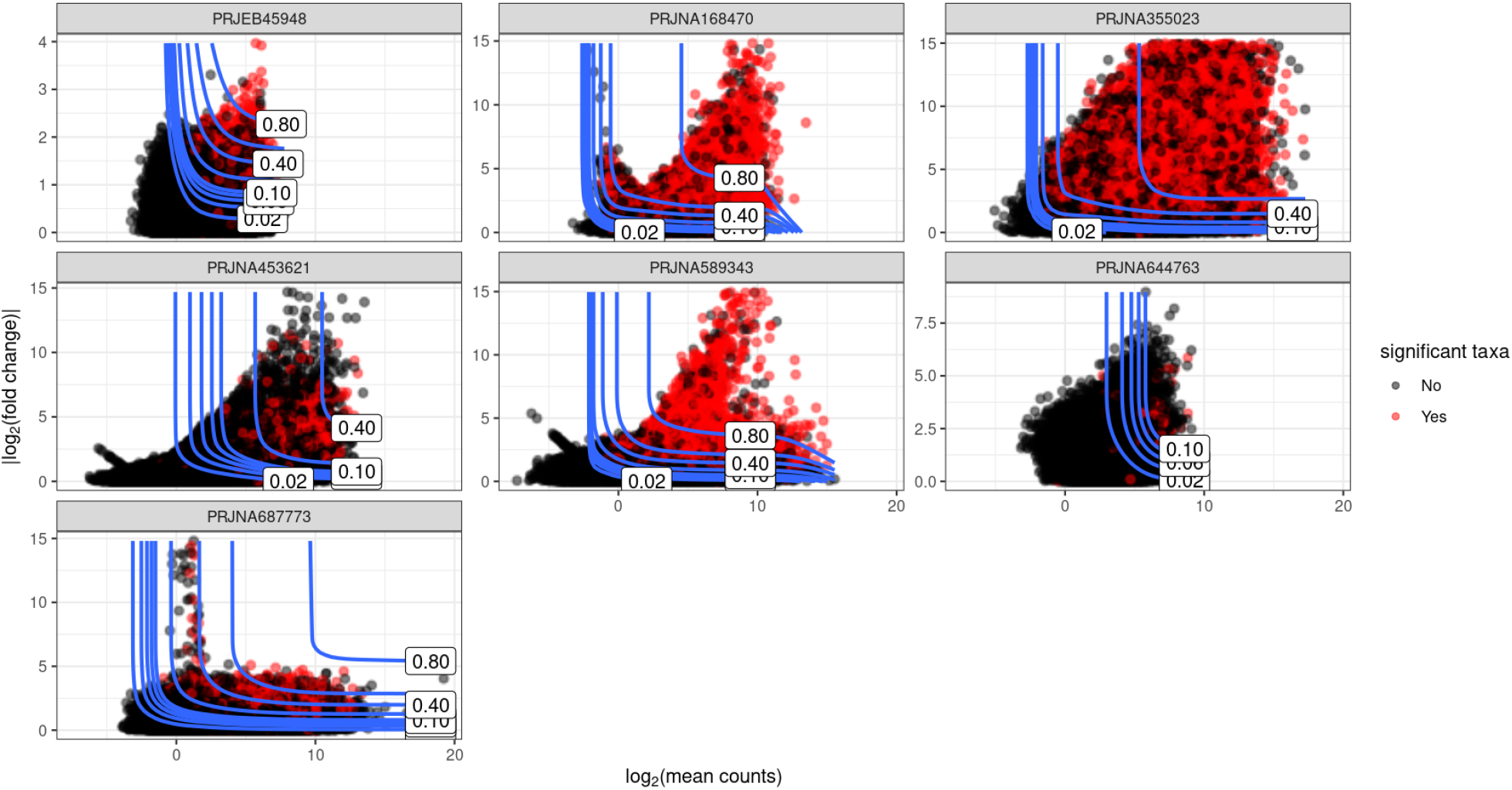
Contour plot showing statistical power for various combinations of overall log mean abundance and log fold change (1000 taxa, 100 samples per group and 100 simulations). Red points indicate simulated taxa with significant log fold change (that is, adjusted FDR-corrected *p*-value *<* 0.1); black points show simulated taxa where we failed to reject the null hypothesis (adjusted *p*-value *>* 0.1). Contour lines show the predicted statistical power for various combinations of log mean abundance and log fold change.

**Fig. 5.**
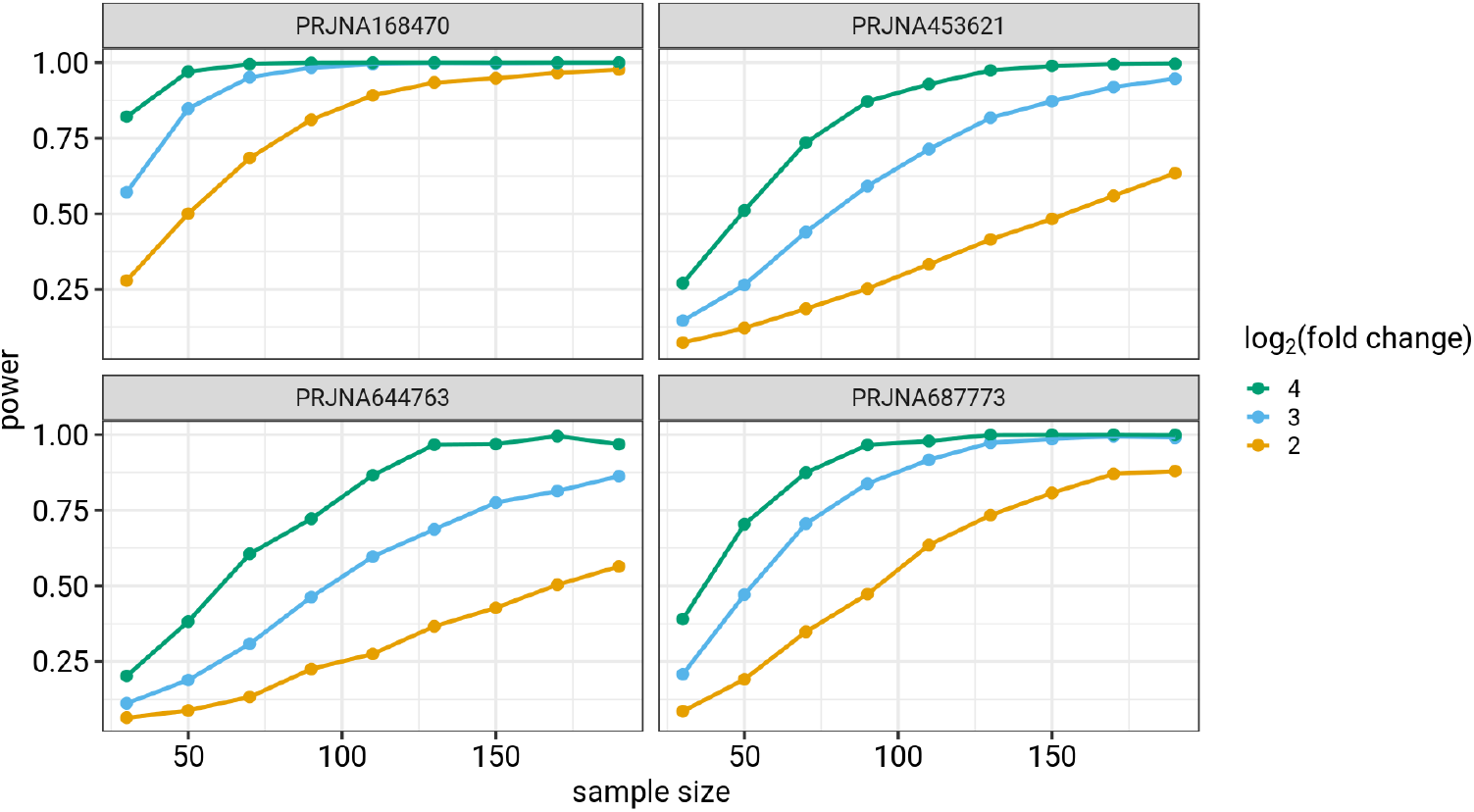
Relationship between statistical power, sample size and log fold change (log mean abundance = 5)

Figure 6 shows the expected number of taxa per experiment with significant differences between groups. Increasing sample size increases the expected number of significant taxa. Figure 7 compares the average power (defined by the arithmetic mean of the power estimates for all taxa) with the quantiles of power estimates for the individual taxon.

**Fig. 6.**
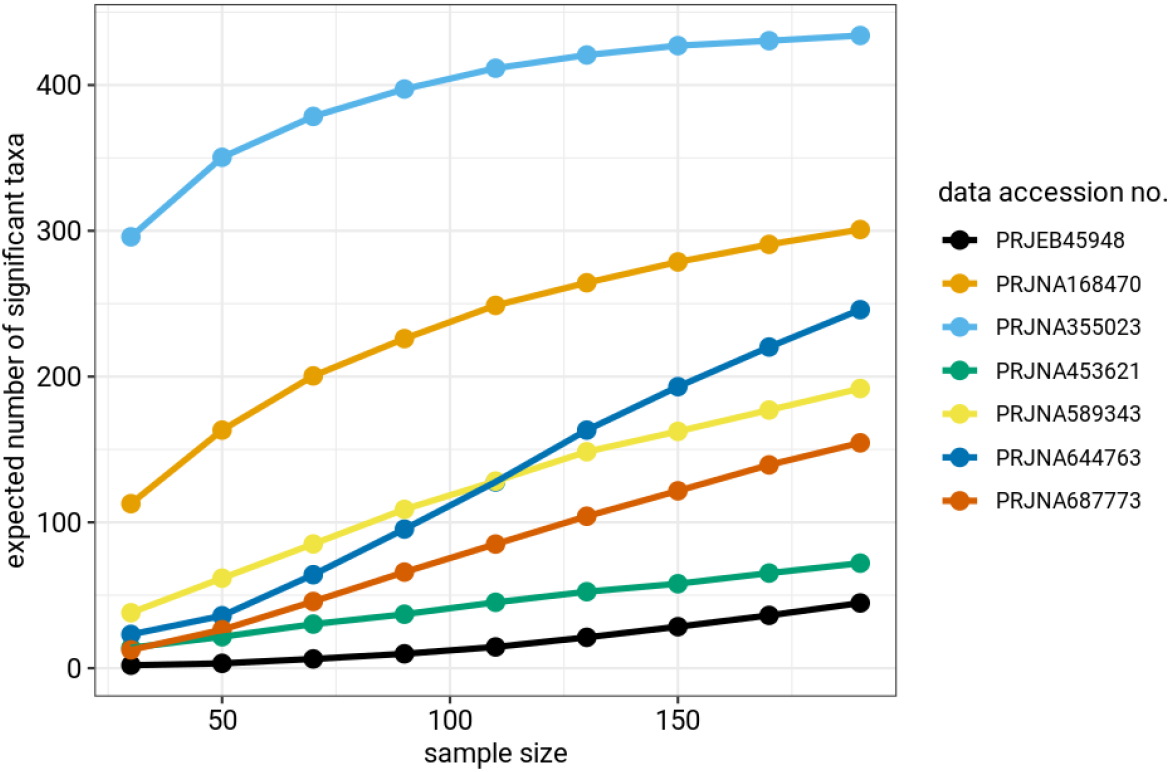
Expected number of significant taxa (out of 1000) for 30, 50, 70, 90, 110, 130, 150, 170 and 190 samples per group.

**Fig. 7.**
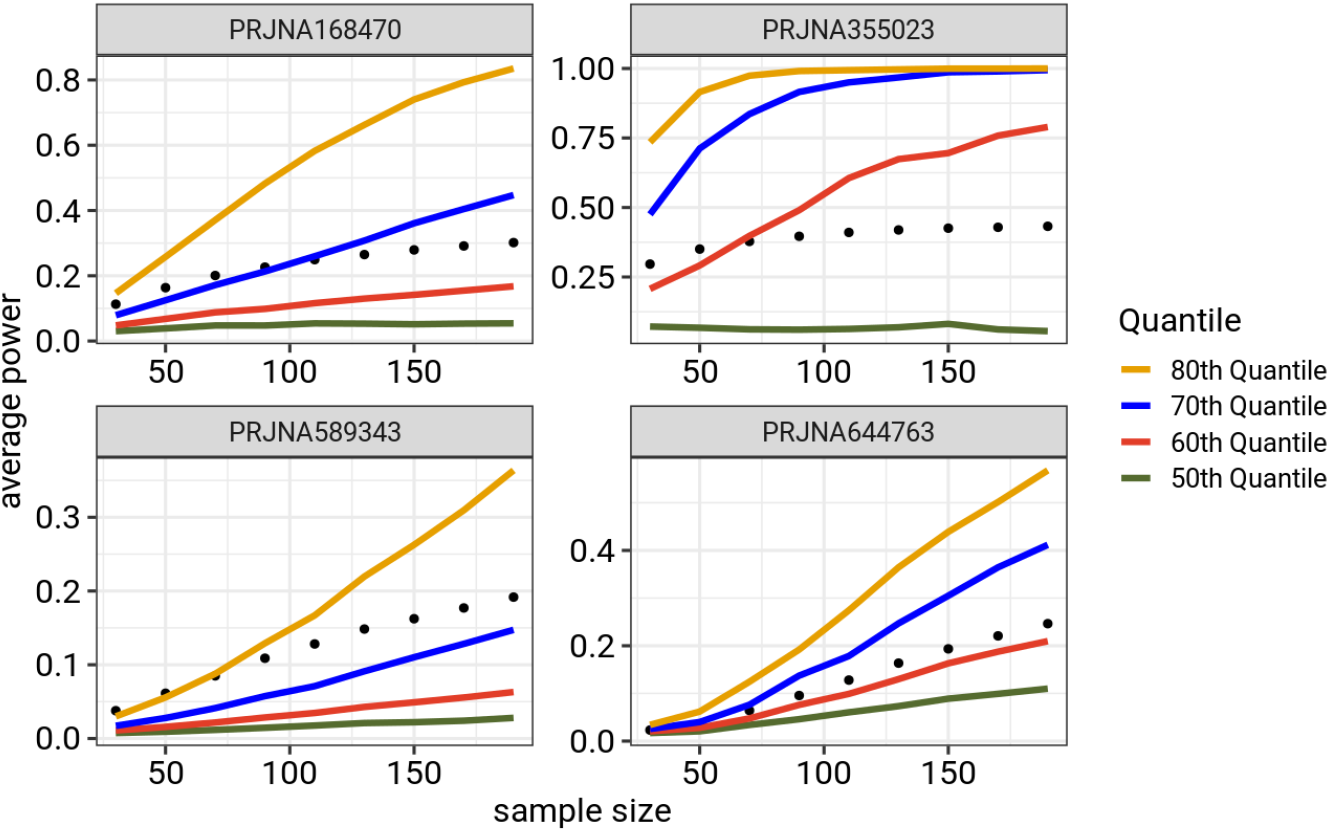
Comparison of average power across power estimates for all taxa and quantiles of taxon-by-taxon power estimates. Average power often overestimates the statistical power for most taxa and might not be a good metric for understanding the power to detect effects in a differential abundance study, hence the need to estimate power at the level of individual taxa.

## Discussion

### Simulated microbiome data

The distributions of mean abundance and variance of taxa from our simulation method match those from the microbiome dataset well (Figs 2 and 3). In contrast, the `HMP` simulations fail to accurately replicate the distributions of taxon means and variance from the data. `metaSPARSim` effectively mimics the mean abundance of taxa but may struggle to replicate the variance distribution.

### Low statistical power

Figure 4 shows a strong positive relationship between statistical power and fold change, as well as a weak positive relationship between mean abundance and statistical power, as anticipated. Few simulated taxa are in regions of high power (80% is the usual target for power in most scientific fields (Cohen, 2013; Descôteaux, 2007)), making it unlikely to attain high power in practical scenarios. Most individual taxa, in most data sets, have power less that 80%.

Our simulation studies were conducted with a relatively large sample sizes (for the field of microbiome studies in health sciences) of 100 samples per group. However, most microbiome studies are often constrained by practical limitations that restrict the available sample size. For example, Kers and Saccenti (2021) examined 100 publications and found a median sample size of 39 samples per group, with a mode of 8 samples. Given the prevalence of low sample sizes in the microbiome literature, differential abundance microbiome studies might have even lower power to detect biologically meaningful effects for individual taxon than those suggested by Figure 4.

The expected number of significant taxa we calculated showed many fewer expected significant taxa for lower sample sizes (Figure 6). This implies that studies with low sample sizes stand the risk of missing taxa with biologically significant effects. Differential abundance microbiome studies might therefore require higher sample sizes than those prevalent in the literature in order to identify majority of the taxa with biologically significant effects. Low statistical power has significant impacts on the reliability of research results. Not only does it lead to type II errors (false negatives) but also causes strong upward bias in the magnitude of estimated effect sizes, via the “winner’s curse” when a statistical significance filter (i.e., taxa with *p*-values below a threshold) is applied (Button et al., 2013).

### Mean power estimate versus taxon by taxon power estimate

Figure 7 shows the relationship between sample size and the average of the power estimates across all taxa. The figure also shows quantiles of power estimates for taxa in each sample size. The average power is needed for computing the expected number of significant taxa (see section 2.6). The average power, however does not provide accurate understanding of the statistical power of the individual taxon in a differential abundance study. The average statistical power for each microbiome dataset is consistently higher than the 50^th^ quantile of the individual taxon-by-taxon power estimates (Figure 7). In most cases, the average power surpasses the 60^th^ quantile of the individual power estimates, indicating that the average power overestimates the power for the majority of taxa. This highlights the need to consider statistical power at the level of individual taxon in a differential abundance study. Average power might overstate the power for most taxa and may lead researchers to underestimate the required sample sizes for their studies.

## Conclusion

Our study sheds light on potentially low statistical power to detect effect size of individual taxa in differential abundance microbiome studies. We introduced a novel method to estimate statistical power for individual taxon. Our method estimates power as a function of fold change and mean abundance of individual taxon. We also introduced a novel simulation method tailored to microbiome count data using seven diverse datasets from ASD children. This approach involved fitting finite Gaussian mixture distributions to estimate key parameters, aligning well with observed data.

Contour plots showing power for individual taxon suggest potentially low power to detect effect size of individual taxon in a differential abundance microbiome study (Figure 4). Low statistical power for individual taxa suggests that differential abundance studies might be missing many taxa with meaningful biological effects (Figure 6). Our findings also show that differential abundance studies may require larger sample sizes than are currently prevalent in microbiome research in order to achieve adequate statistical power (Figure 5).

The power estimation method presented in this study will enable researchers to estimate power at the level of individual taxon, quantify the range of power across all taxa, and estimate the expected number of significant taxa for their study. Our framework and simulation-based evidence contribute to enhancing understanding in the field, promoting accurate result interpretation. The provided framework and code facilitate reproducibility and empower researchers to make informed decisions about study design.

## Author contributions statement

B.B provided the initial concept. M.A conducted all analyses and wrote the first draft of the manuscript. B.B and M.A reviewed the analyses and wrote the final manuscript together.

## Acknowledgments

This work is supported in part by the Natural Sciences and Engineering Research Council (NSERC) Discovery grants (NSERC # 2016-05488 and # 2023-05400).

## Supplementary Material

**Fig. 8.**
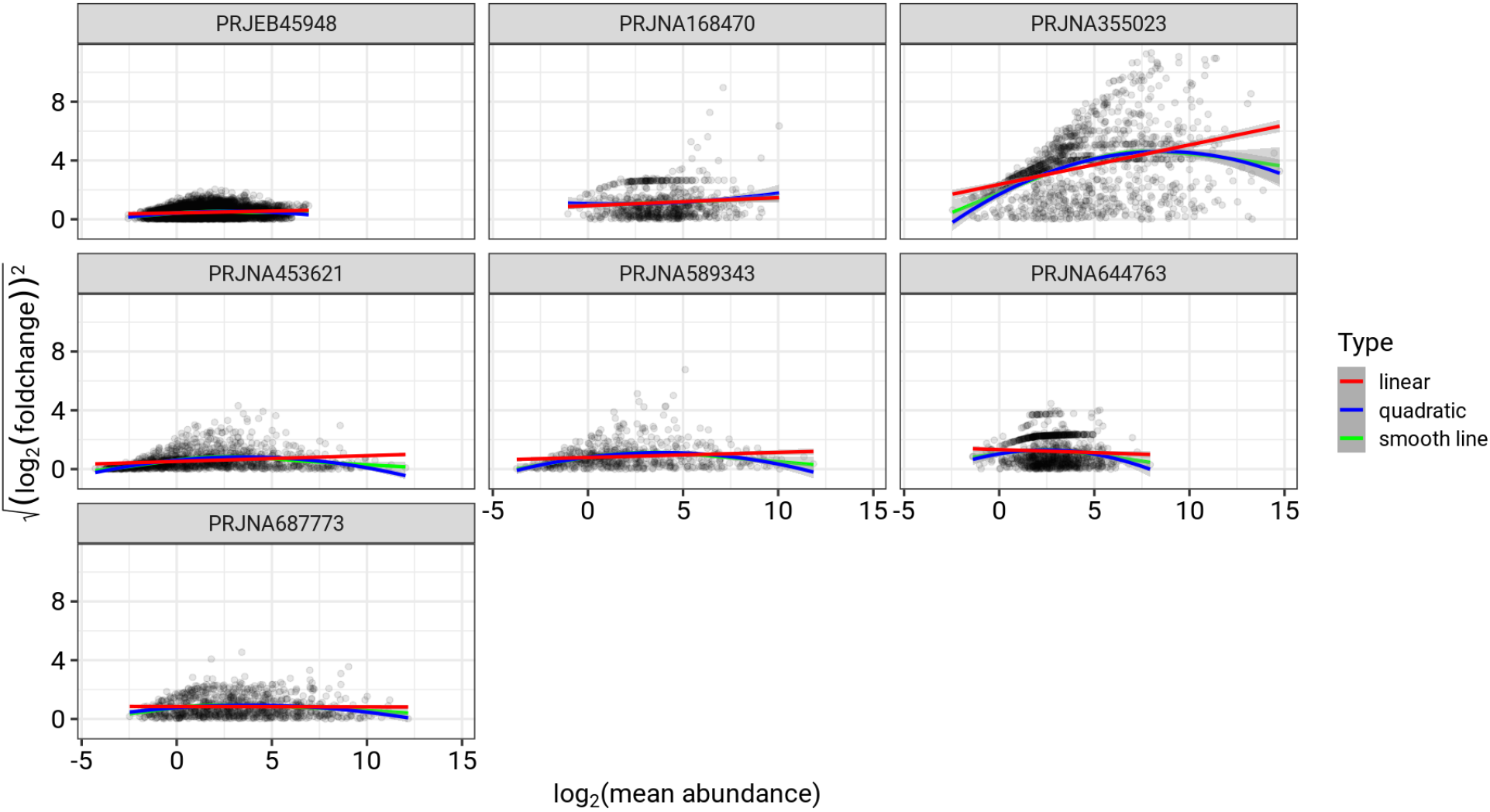
Scale location plots to determine functions to model standard deviation parameter

**Fig. 9.**
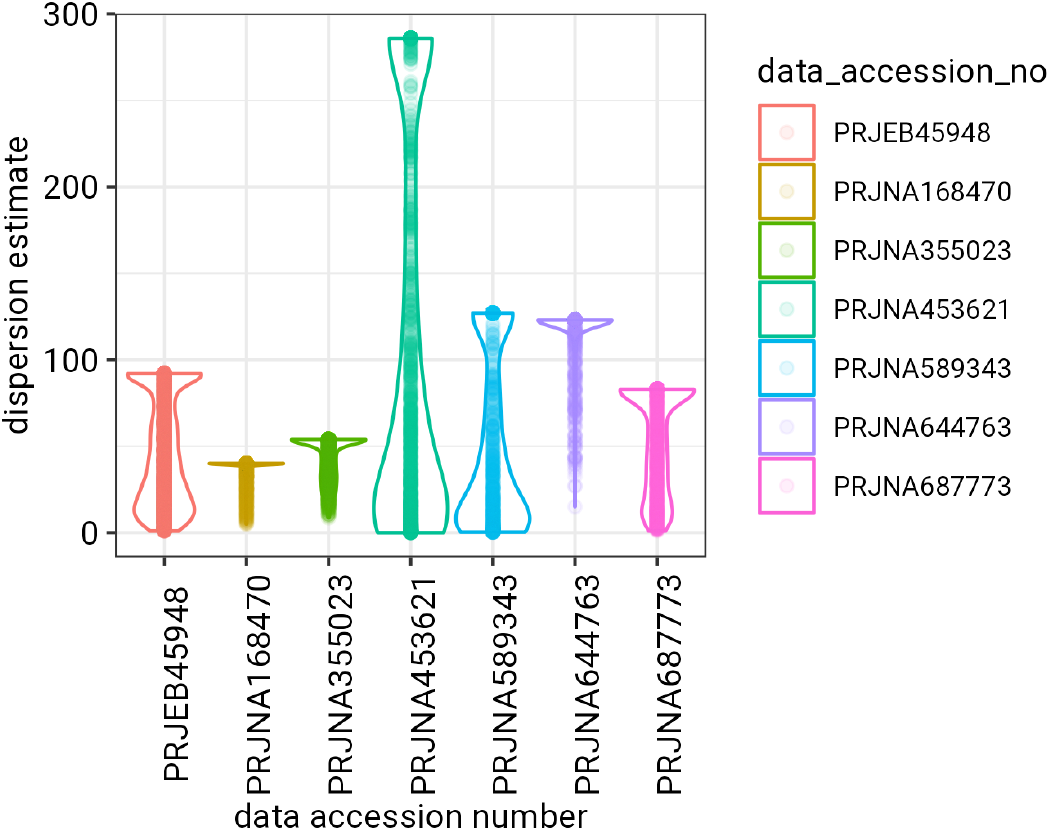
Distribution of dispersion estimates from DESeq2

**Fig. 10.**
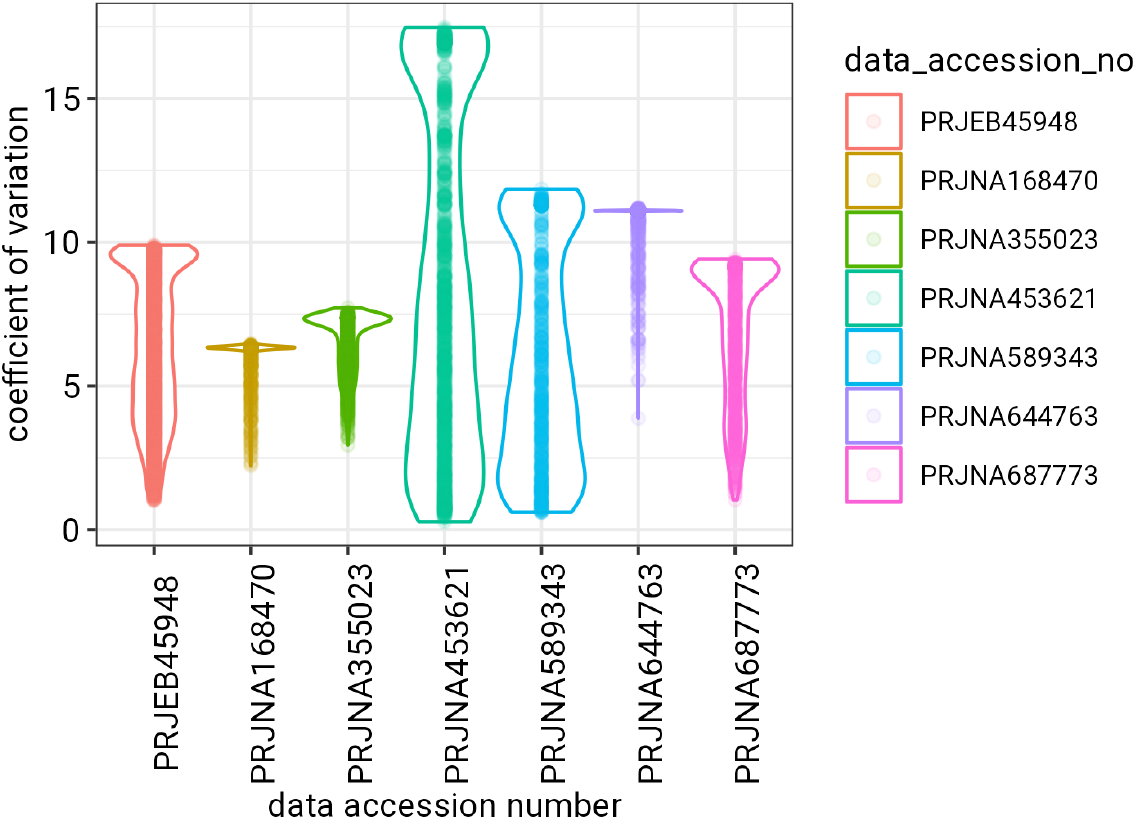
Coefficient of variation of taxa abundance using dispersion estimates from DESeq2

